# CD8^+^ T cell immunogenicity induced by endogenous EVs engineered by antigens fused to a truncated Nef^mut^ EV-anchoring protein

**DOI:** 10.1101/2021.02.05.429891

**Authors:** Chiara Chiozzini, Francesco Manfredi, Flavia Ferrantelli, Patrizia Leone, Andrea Giovannelli, Eleonora Olivetta, Maurizio Federico

## Abstract

Intramuscular injection of DNA vectors expressing the extracellular vesicle (EV)-anchoring protein Nef^mut^ fused at its C-terminus to viral and tumor antigens elicits a potent, effective, and anti-tolerogenic CD8^+^ T cell immunity against the heterologous antigen. The immune response is induced through the production of EVs incorporating Nef^mut^-derivatives released by muscle cells. In the perspective to a possible translation into the clinic of the Nef^mut^-based vaccine platform, we aimed at increasing its safety profile by identifying the minimal part of Nef^mut^ retaining the EV-anchoring protein property. We found that a C-terminal deletion of 29-amino acids did not affect the ability of Nef^mut^ to associate with EVs. Furthermore, the EV-anchoring function was preserved when antigens from both HPV16 (i.e., E6 and E7) and SARS-CoV-2 (i.e., S1 and S2) were fused to its C-terminus. By analyzing the immune responses induced after intramuscular injection of DNA vectors expressing fusion products based on the four viral antigens, we found that the Nef^mut^ C-terminal deletion did not impact on the levels of antigen –specific CD8^+^ T lymphocytes as evaluated by IFN-γ EliSpot analysis and intracellular cytokine staining. In addition, immune responses at distal sites remained unaffected, as indicated by the similar percentages of SARS-CoV-2 S1- and S2-specific CD8^+^ T cells detected in spleens and lung airways of mice injected with DNA vectors expressing the viral antigens fused with either Nef^mut^ or Nef^mut^PL.

We concluded that the C-terminal Nef^mut^ truncation does not affect stability, EV-anchoring, and CD8^+^ T cell immunogenicity of the fused antigen. Hence, Nef^mut^PL represents a safer alternative to full-length Nef^mut^ for the design of CD8^+^ T cell vaccines for humans.

## Introduction

Extracellular vesicles (EVs) are classified as apoptotic bodies (1-5 µm), microvesicles (50-1,000 nm), and exosomes (50-200 nm) (1). Microvesicles (also referred to as ectosomes) shed by plasma membrane, whereas exosomes are released after inward invagination of endosome membranes and formation of intraluminal vesicles. Exosomes are secreted by all cell types and are an important mean of intercellular communication by transporting their cargo, such as DNAs, RNAs, proteins, and lipids, from the producer cell to the recipient one (2). Healthy cells constitutively release both exosomes and microvesicles, together referred to as EVs hereinafter.

Nef^mut^ is a Human Immunodeficiency Virus-1 protein mutant defective for all anti-cellular Nef functions including CD4 and MHC Class I down-regulation, increased HIV-1 infectivity, and p21 activated kinases (PAK)-2 activation (3). We previously described the high efficiency of HIV-1 Nef^mut^ of uploading into EVs released by multiple cell types (4). The efficiency of Nef^mut^ incorporation into EVs is maintained also when a foreign protein is fused to its C-terminus (5-7). When DNA vectors expressing Nef^mut^-based fusion proteins are intramuscularly (i.m.) injected in mice, significant quantities of the fusion proteins are packed into EVs while not altering their spontaneous release from muscle tissue. These *in vivo* engineered EVs can freely circulate into the body, and internalized by antigen-presenting cells (APCs) which cross-present EV contents to activate antigen-specific CD8^+^ T cells. These events result in the induction of potent antigen-specific CD8^+^ T cytotoxic lymphocyte (CTL) responses (6-8). Effectiveness and flexibility of this vaccine platform have been demonstrated with an array of viral products of various origins and sizes including but not limited to: Human Papilloma virus (HPV)16-E6 and -E7; Severe acute Respiratory Syndrome Coronavirus (SARS-CoV)-2 S1, S2, M, and N; Ebola Virus VP24, VP40 and NP; Hepatitis C Virus NS3; West Nile Virus NS3; and Crimean-Congo Hemorrhagic Fever NP (7).

To maximize the safety profile of the Nef^mut^-based immunogens, we tried to identify the minimum part of Nef^mut^ still retaining in full the efficiency of EV-anchoring protein and the associated EV immunogenicity. Investigations were carried out in the context of two viral systems recognizing quite different pathogenic effects, i.e., HPV-16, whose infection can induce tumors, and the acute respiratory disease-inducing SARS-CoV-2.

Pre-clinical studies already provided evidence that the Nef^mut^-based CTL vaccine platform can be exploited as therapeutic intervention against HPV16-related and other malignancies (6, 9). The benefits expected from a therapeutic cancer vaccine refer to the possibility to induce a *de novo* antitumor immunity as well as widen both potency and breadth of pre-existing immunity (10). Cancer cells can express a multitude of new antigens as result of their intrinsic genetic instability typical of malignant transformation and/or of the expression of the etiologic cancer agents, as in the case of virus-induced malignancies, e.g., HPV and HBV. In non-virus-induced cancers, transformed cells can produce antigens to which the host is basically tolerant (tumor-associated self-antigens) as in the case of antigens expressed during fetal development (oncofetal antigens), and overexpressed in specific tissues, e.g., HER2 in mammary glands. In addition, cancer cells can express antigens to which the host does not develop tolerance, being however the immune response is not effective enough to counteract the cancer cell growth. They include the so-called “tumor specific neo-antigens” as well as antigens normally produced in immune-privileged tissues, as for cancer-testis antigens (11). Hence, establishing a safe method to induce an adaptive immune response against both tolerogenic and non-tolerogenic TAAs would be of great relevance for the design of novel antitumor therapeutic approaches.

Previous work demonstrated the role of specific CD8^+^ T cell immunity in attenuating the disease induced by influenza virus (12, 13). Similarly, both experimental and clinical evidences supported the idea that a SARS-CoV-2-specific CD8^+^ T cell immunity can be instrumental to at least strongly mitigate the symptoms related to the viral spread in both upper and lower airways (14-16). Data from experimental infections in rhesus macaques indicated that the virus-specific CD8^+^ T cell immunity is critical to protect the animals from virus re-challenge after the rapid decay of neutralizing antibodies (17). Consistently, induction of antiviral CD8^+^ T cells was associated with a strongly reduced severity of the disease in humans (18). Furthermore, the demonstrated ability of SARS-CoV-2 to spread through cell-to-cell contact (19) implies the necessity to induce a robust cellular immunity to contain and clear the virus. In this context, the development of novel preventive strategies focused on the induction of anti-SARS-CoV-2 CD8^+^ T cell immunity would be considered of relevance.

The Nef^mut^-based CTL vaccine platform has been already proven to be a candidate for both antiviral and antitumor vaccines. In this perspective, optimization of the safety profile of Nef^mut^-based immunogens is mandatory. This aim can be pursued, for instance, by removing unnecessary sequences from the anchoring protein. Here, we provide evidence that both EV-incorporation efficiency and immunogenicity of foreign antigens are conserved in the presence of a C-terminal 29 amino acid deletion of Nef^mut^.

## Materials and Methods

### DNA vector synthesis

The pTargeT (Invitrogen, Thermo Fisher Scientific) Nef^mut^ and Nef^mut^fusion vectors were already described (5, 6). The pTargeT-Nef^mut^/E6 vector comprised an E6 ORF which was codon optimized through an *ad hoc* algorithm provided by Codon Optimization On-Line (COOL) service (https://cool.syncti.org) which introduced 134 base substitutions. In addition, the ^G^130^V^ amino acid substitution generated a loss-of-function of E6 by hindering its interaction with the p53 cell protein partner (20). The E6 ORF was inserted in *Apa* I/*Sal* I sites of the pTargeT-Nef^mut^fusion vector. The pTargeT-Nef^mut^/E7 vector was obtained through a similar strategy. The E7 ORF was codon optimized through the introduction of 64 base substitutions as described by Cid-Arregui and coll. (21). The protein was detoxified through the insertion of three amino acid substitutions, namely three glycines at positions 21, 24, and 26, within the retinoblastoma protein (pRB) binding site. In this way, the E7-specific immortalizing activity was abrogated (21).

The pTargeT-Nef^mut^PL vector was obtained starting from the vector pTargeT-Nef^mut^. It was digested with *Sma* I which cuts just downstream to the most C-terminal typical Nef^mut^ mutation (i.e., ^E^177^G^), and at the 3’ end of vector polylinker. The subsequent re-ligation generated a C-terminal 29 amino acid deletion, with a *de novo* formation of a stop codon just downstream the *Sma* I restriction site.

To obtain the pTargeT Nef^mut^-PL-based DNA vectors, an intermediate construct referred to as Nef^mut^PLfusion was obtained. To this aim, the Nef^mut^PL ORF from pTargeT-Nef^mut^PL was PCR amplified using a forward primer tagged with a *Nhe* I restriction site, and a reverse primer including an *Apa* I site as well as the overlapping sequence for a GPGP linker. The PCR product was then inserted in the corresponding sites of pTargeT vector. In this way, the insertion in the unique *Apa* I restriction site of downstream ORFs resulted in an in-frame sequence. To obtain the pTarget-Nef^mut^PL/E6 and pTarget-Nef^mut^PL/E7 vectors, synthesis and cloning strategy were identical to that described for the DNA vectors expressing the full-length Nef^mut^.

Both Nef^mut^/SARS-CoV-2-based fusion proteins were cloned into the pVAX1 plasmid (Thermo Fisher) as already described (22). Both pVAX1-Nef^mut^ and pVAX1-Nef^mut^PL vectors were obtained by inserting the respective ORFs in *Nhe* I and *Eco R*I sites of the vector polylinker. To obtain pVAX1 vectors expressing Nef^mut^PL fused with either SARS-CoV-2 S1 and S2 ORFs, an intermediate construct referred to as pVAX1-Nef^mut^PLfusion was obtained. To this aim, the Nef^mut^PL ORF from pTargeT-Nef^mut^PL was PCR amplified using a forward primer tagged with a *Nhe* I restriction site, and a reverse primer including an *Apa* I as well as the overlapping sequence for a GPGP linker, and inserted in the corresponding sites of pVAX1 vector. In this way, the downstream insertion of S1 and S2 ORFs in *Apa* I and *Pme* I restriction sites resulted in in-frame sequences, and, upon translation, in Nef^mut^PL-based fusion proteins. Gene synthesis were carried out by Eurofins Genomics and Explora Biotech.

### Cell cultures and transfection

Human embryonic kidney (HEK)293T cells (ATCC, CRL-11268) were grown in DMEM (Gibco) plus 10% heat-inactivated fetal calf serum (FCS, Gibco). Transfection assays were performed using Lipofectamine 2000 (Invitrogen, Thermo Fisher Scientific).

### EV isolation

Cells transfected with vectors expressing the Nef^mut^-based fusion proteins were washed 24 h later, and reseeded in medium supplemented with EV-deprived FCS. The supernatants were harvested from 48 to 72 h after transfection. EVs were recovered through differential centrifugations (23) by centrifuging supernatants at 500×*g* for 10 min, and then at 10,000×*g* for 30 min. Supernatants were harvested, filtered with 0.22 µm pore size filters, and ultracentrifuged at 70,000×*g* for l h. Pelleted vesicles were resuspended in 1×PBS, and ultracentrifuged again at 70,000×*g* for 1 h. Afterwards, pellets containing EVs were resuspended in 1:100 of the initial volume.

### Western blot analysis

Western blot analyses of both cell lysates and EVs were carried out after resolving samples in 10% sodium dodecyl sulfate-polyacrylamide gel electrophoresis (SDS-PAGE). In brief, the analysis on cell lysates was performed by washing cells twice with 1×PBS (pH 7.4) and lysing them with 1x SDS-PAGE sample buffer. Samples were resolved by SDS-PAGE and transferred by electroblotting on a 0.45 μM pore size nitrocellulose membrane (Amersham) overnight using a Bio-Rad Trans-Blot. For western blot analysis of EVs, they were lysed and analyzed as described for cell lysates. For immunoassays, membranes were blocked with 5% non-fat dry milk in PBS containing 0.1% Triton X-100 for 1 h at room temperature, then incubated overnight at 4 °C with specific antibodies diluted in PBS containing 0.1% Triton X-100. Filters were revealed using 1:1,000-diluted sheep anti-Nef antiserum ARP 444 (MHRC, London, UK), 1:500-diluted anti-β-actin AC-74 mAb from Sigma, and 1:500 diluted anti-Alix H-270 polyclonal Abs from Santa Cruz.

### Mice immunization

Both 6-weeks old C57 Bl/6 and, for Nef^mut^/S2 immunizations (in view of the lack of already characterized H2^b^ immunodominant S2 epitopes), Balb/c female mice were obtained from Charles River. They were hosted at the Central Animal Facility of the Istituto Superiore di Sanità (ISS), as approved by the Italian Ministry of Health, authorization n. 565/2020 released on June, 3, 2020. Preparations of DNA vector were diluted in 30 µL of sterile 0.9% saline solution. Both quality and quantity of the DNA preparations were checked by 260/280 nm absorbance and electrophoresis assays. Each inoculum volume was injected into both quadriceps. Mice were anesthetized with isoflurane as prescribed in the Ministry authorization.. Immediately after inoculation, mice underwent electroporation at the site of injection through the Agilpulse BTX device using a 4-needle array 4 mm gap, 5 mm needle length, with the following parameters: 1 pulse of 450 V for 50 µsec; 0.2 msec interval; 1 pulse of 450 V for 50 µsec; 50 msec interval; 8 pulses of 110 V for 10 msec with 20 msec intervals. The same procedure was repeated for both quadriceps of each mouse. Immunizations were repeated after 14 days. Fourteen days after the second immunization, mice were sacrificed by either cervical dislocation or CO_2_ inhalation.

### Cell isolation from immunized mice

Spleens were explanted by qualified personnel of the ISS Central Animal Facility, and placed into a 2 mL Eppendorf tubes filled with 1 mL of RPMI 1640 (Gibco), 50 µM 2-mercaptoethanol (Sigma). Spleens were transferred into a 60 mm Petri dish containing 2 mL of RPMI 1640 (Gibco), 50 µM 2-mercaptoethanol (Sigma). Splenocytes were extracted by incising the spleen with sterile scissors and pressing the cells out of the spleen sac with the plunger seal of a 1 mL syringe. After addition of 2 mL of RPMI medium, cells were transferred into a 15 mL conical tube and the Petri plate was washed with 4 mL of medium to collect the remaining cells. After a three-minute sedimentation, splenocytes were transferred to a new sterile tube to remove cell/tissue debris. Counts of live cells were carried out by the trypan blue exclusion method. Fresh splenocytes was resuspended in RPMI complete medium, containing 50 µM 2-mercaptoethanol and 10% FBS, and tested by IFN-γ EliSpot assay.

For bronchoalveolar lavages, mice were sacrificed by CO_2_ inhalation, placed on their back, and dampened with 70% ethanol. Neck skin was opened to the muscles by scissors, and muscles around the neck and salivary glands were gently pulled aside with forceps to expose the trachea. A 15 cm long surgical thread was then placed around the trachea and a small hole was cut by fine point scissors between two cartilage rings. A 22 G × 1” Exel Safelet Catheter was carefully inserted about 0.5 cm into the hole, and then the catheter and the trachea were firmly tied together with the suture. A 1 mL syringe was loaded with 1 mL of cold PBS and connected to the catheter. The buffered solution was gently injected into the lung and aspirated while massaging mouse torax. Lavage fluid was tranferred to a 15 mL conical tube on ice, and lavage repeated two more times (24, 25). Total lavage volume was approximately 2.5 mL/mouse. Cells were recovered by centrifugation, resuspeded in cell culture medium, and counted.

### IFN-γ EliSpot analysis

A total of 2.5×10^5^ live cells were seeded in each microwell. Cultures were run in triplicate in EliSpot multiwell plates (Millipore, cat n. MSPS4510) pre-coated with the AN18 mAb against mouse IFN-γ (Mabtech) in RPMI 1640 (Gibco), 10% FCS, 50 µM 2-mercaptoethanol (Sigma) for 16 h in the presence of 5 µg/mL of the following CD8-specific peptides: E6 (H2-K^b^): 18-26: KLPQLCTEL (27); 50-57: YDFAFRDL (27); 109-117: RCINCQKPL (28); 127-135: DKKQRFMNI (27). HPV-16 E7 (H2-K^b^): 49-57 RAHYNIVTF (27); 67-75 LCVQSTHVD (28). SARS-CoV-2 S1 (H2-K^b^): 525-531 VNFNFNGL (29); SARS-CoV-2 S2 (H2-K^d^): 1079-1089 PAICHDGKAH (30). As a negative control, 5 µg/mL of either H2-K^b^ or H2-K^d^-binding peptides were used. More than 70% pure preparations of the peptides were obtained from both UFPeptides, Ferrara, Italy, and JPT, Berlin, Germany. For cell activation control, cultures were treated with 10 ng/mL PMA (Sigma) plus 500 ng/mL of ionomycin (Sigma). After 16 hours. cultures were removed, and wells incubated with 100 µL of 1 µg/ml of the R4-6A2 biotinylated anti-IFN-γ (Mabtech) for 2 hours at r.t. Wells were then washed and treated for 1 hour at r.t. with 1:1000 diluted streptavidine-ALP preparations from Mabtech. After washing, spots were developed by adding 100 µL/well of SigmaFast BCIP/NBT, Cat. N. B5655. The spot-forming cells were finally analyzed and counted using an AELVIS EliSpot reader.

### Intracellular cytokine staining (ICS)

Splenocytes were seeded at 2×10^6^/mL in RPMI medium, 10% FCS, 50 µM 2-mercaptoethanol (Sigma), and 1 µg/mL brefeldin A (BD Biosciences). Control conditions were carried out either by adding 10 ng/ml PMA (Sigma) and 1 µg/mL ionomycin (Sigma), or with unrelated peptides. After 16 hours, cultures were stained with 1 µl of LIVE/DEAD Fixable Aqua Dead Cell reagent (Invitrogen ThermoFisher) in 1 mL of PBS for 30 minutes at 4 °C and washed twice with 500 µl of PBS. To minimize nonspecific staining, cells were pre-incubated with 0.5 µg of Fc blocking mAbs (i.e., anti-CD16/CD32 antibodies, Invitrogen/eBioscience) in 100 µL of PBS with 2% FCS for 15 minutes at 4 °C. For the detection of cell surface markers, cells were stained with 2 µL of the following Abs: FITC conjugated anti-mouse CD3, APC-Cy7 conjugated anti-mouse CD8a, and PerCP conjugated anti-mouse CD4 (BD Biosciences) and incubated for 1 hour at 4 °C. After washing, cells were permeabilized and fixed through the Cytofix/Cytoperm kit (BD Biosciences) as per the manufacturer’s recommendations, and stained for 1 hour at 4 °C with 2 µl of the following Abs: PE-Cy7 conjugated anti-mouse IFN-γ (BD Biosciences), PE conjugated anti-mouse IL-2 (Invitrogen eBioscience), and BV421 anti-mouse TNF-α (BD Biosciences) in a total of 100 µL of 1× Perm/Wash Buffer (BD Biosciences). After two washes, cells were fixed in 200 µL of 1× PBS/formaldehyde (2% v/v). Samples were then assessed by a Gallios flow cytometer and analyzed using Kaluza software (Beckman Coulter).

Gating strategy was as follows: live cells as detected by Aqua LIVE/DEAD Dye vs. FSC-A, singlet cells from FSC-A vs. FSC-H (singlet 1) and SSC-A vs SSC-W (singlet 2), CD3 positive cells from CD3 (FITC) vs. SSC-A, CD8 or CD4 positive cells from CD8 (APC-Cy7) vs. CD4 (PerCP). The CD8^+^ cell population was gated against APC-Cy7, PE, and BV421 to observe changes in IFN-γ, IL-2, and TNF-□ production, respectively. Boolean gates were created in order to determine any cytokine co-expression pattern.

### Statistical analysis

When appropriate, data are presented as mean + standard deviation (SD). In some instances, the Mann-Whitney U test was used. *p*<0.05 was considered significant.

## Results

### Design and EV-uploading of a C-terminal truncated Nef^mut^

The reduction of sequences of the EV-anchoring protein is expected to increase the safety profile of the Nef^mut^ CTL vaccine platform. We tried to identify the shortest Nef^mut^-related amino acid sequence still retaining the ability to efficiently incorporate into EVs.

HIV-1 Nef is a 206 to 210 amino acid long protein recognizing a quite extended structured N-moiety, and an unstructured C-terminal flexible region (31). To leave untouched the Nef^mut^ secondary structure which was assumed to be necessary for the EV-uploading activity, only isoforms deleted within its unstructured C-terminus were considered. Another relevant constraint was represented by the most C-terminal unique amino acid mutation of Nef^mut^ which maps at position 177, and whose presence was proven to be mandatory to preserve its high efficiency of EV uploading. On this basis, a Nef^mut^ protein truncated at the amino acid 178, hence deprived of 29 C-terminal amino acids, was considered (Fig. 1).

**Figure 1.**
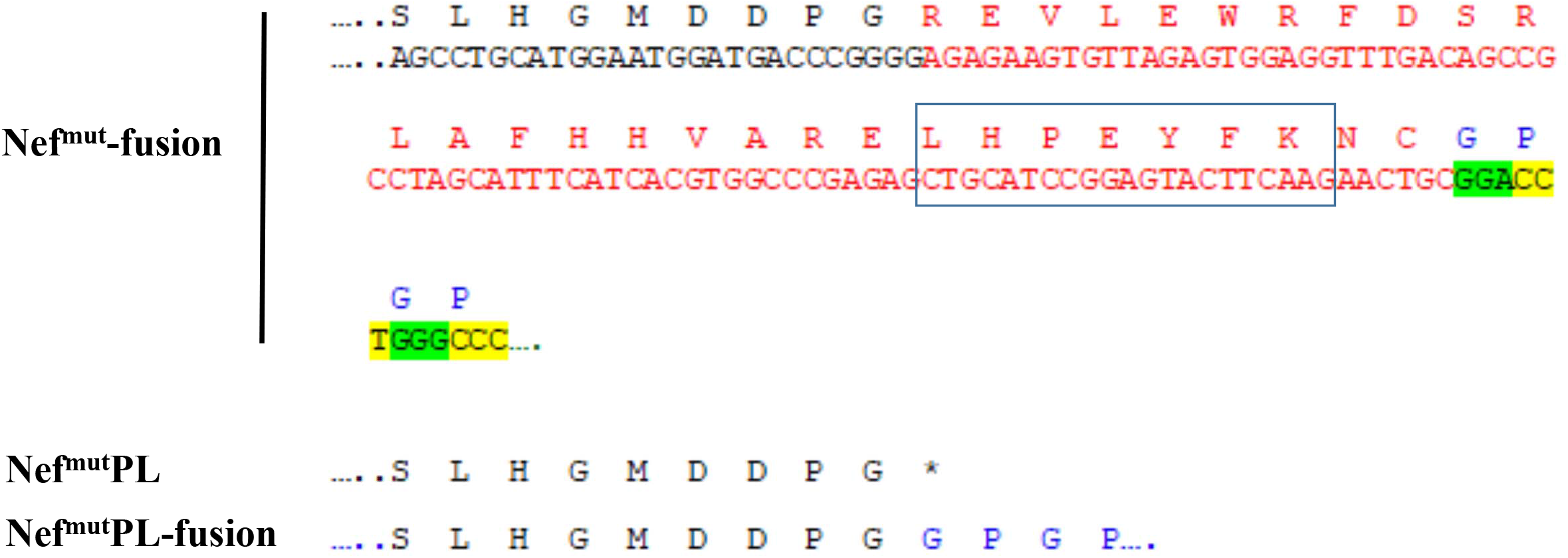
Nef C-terminal sequences in both Nef^mut^PL and Nef^mut^PLfusion DNA expressing vectors. On the top, both nucleotide and amino acid sequences at C-terminus of Nef^mut^ ORF in the context of both pTargeT- and pVAX1-Nef^mut^fusion vectors are reported. Sequences of the 29-amino acid region deleted in the Nef^mut^PL ORF are highlighted in red, and the linker sequence in green/yellow. CRM domain also is highlighted. At the bottom, amino acid sequences of Nef C-terminal regions in both pTargeT-Nef^mut^PL and pTargeT-Nef^mut^PLfusion vectors are reported. In the latter case, the position of the GPGP linker is indicated.

Both cell accumulation and EV association of C-terminal truncated Nef^mut^ (hereinafter referred to as Nef^mut^PL) compared to the full-length isoform were analyzed. To this aim, western blot analysis on lysates of both transiently transfected HEK293T cells and EVs isolated from respective supernatants were carried out. Representative results shown in fig. 2 indicated that Nef^mut^ and Nef^mut^PL accumulated into transfected cells at similar extents, suggesting that the C-terminal truncation did not affect the protein stability. Most important, Nef^mut^PL associated with EVs at levels comparable to those of full-length Nef^mut^, indicating that the presence of the 29 C-terminal amino acids is not essential for the uploading efficiency of Nef^mut^ into EVs.

**Figure 2.**
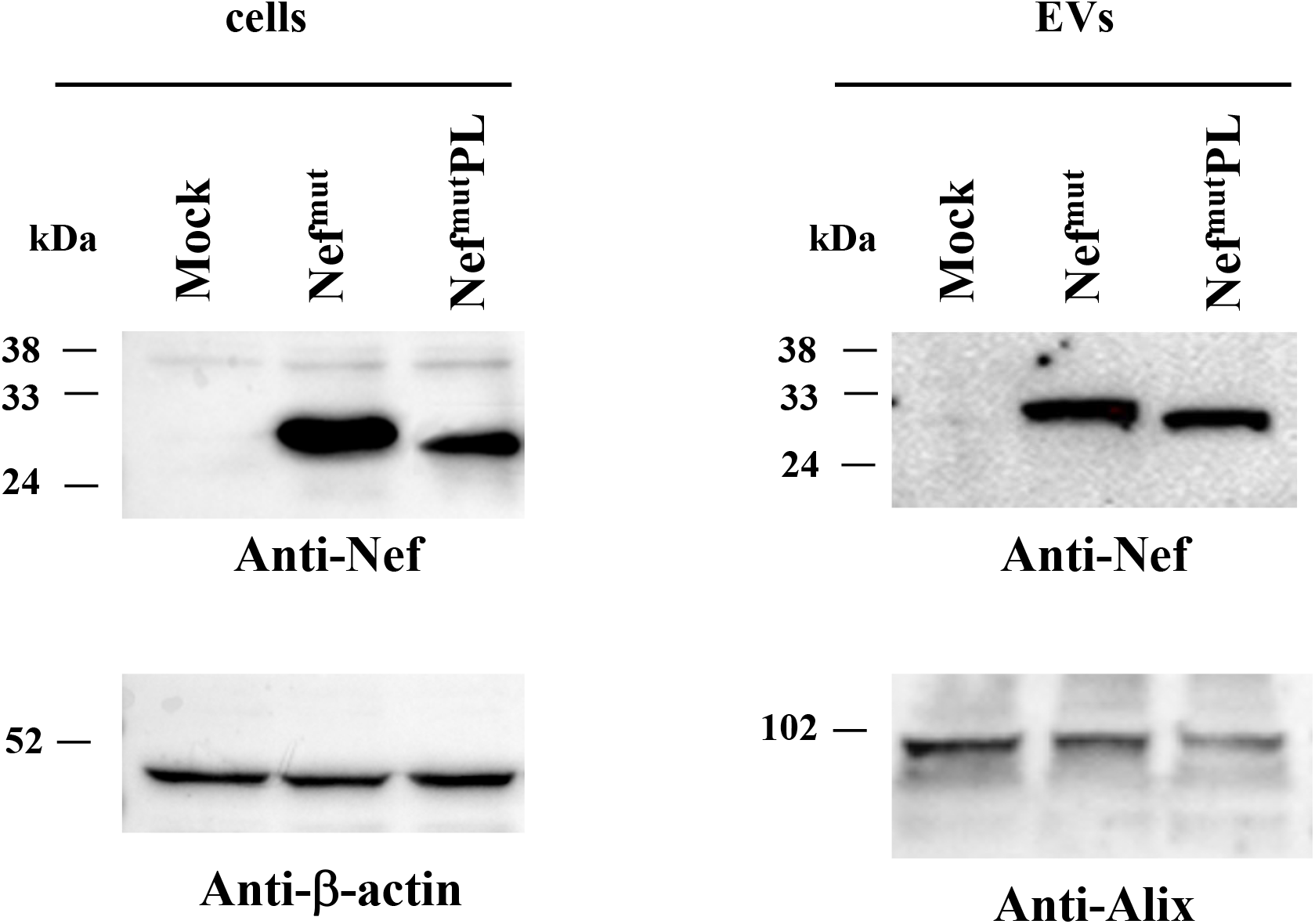
Detection of Nef^mut^PL in transfected cells and EVs. Western blot analysis on 30 µg of lysates of HEK293T cells transfected with DNA vectors expressing either Nef^mut^ or Nef^mut^PL (left panels). Equal volumes of buffer where purified EVs were resuspended after differential centrifugations of the respective supernatants were also analyzed (right panels). As controls, both cell and EV lysates from mock-transfected cells were included. Polyclonal anti-Nef Abs served to detect Nef^mut^-based products, while β-actin and Alix were revealed as markers for cell lysates and EVs, respectively. Molecular markers are given in kDa. The results are representative of seven independent experiments.

### Intracellular expression and EV-uploading HPV16-E6 and -E7 fused with Nef^mut^PL

Next, we tested the efficiency of EV-uploading of foreign proteins fused to Nef^mut^PL compared to parental Nef^mut^. To this aim, we considered both HPV16-E6 and –E7, which were already shown to be uploaded in EVs upon fusion with full-length Nef^mut^ (7, 32). The Nef^mut^-related sequences were fused with either HPV16-E6 or -E7 ORFs which were optimized for translation in eukaryotic cells, and whose protein domains involved in the respective pathogenic effects were inactivated. Steady-state levels in both transfected cells and EVs isolated from the respective supernatants were evaluated by western blot analysis. The E6-based fusion products accumulated in HEK293T transfected cells at similar levels, and the analysis of EVs revealed that E6 efficiently associated with EVs also when fused with Nef^mut^PL (Fig. 3A). Consistently, E7 derivatives fused with either Nef^mut^ or Nef^mut^PL were expressed and uploaded in EVs at similar extents (Fig. 3B).

**Figure 3.**
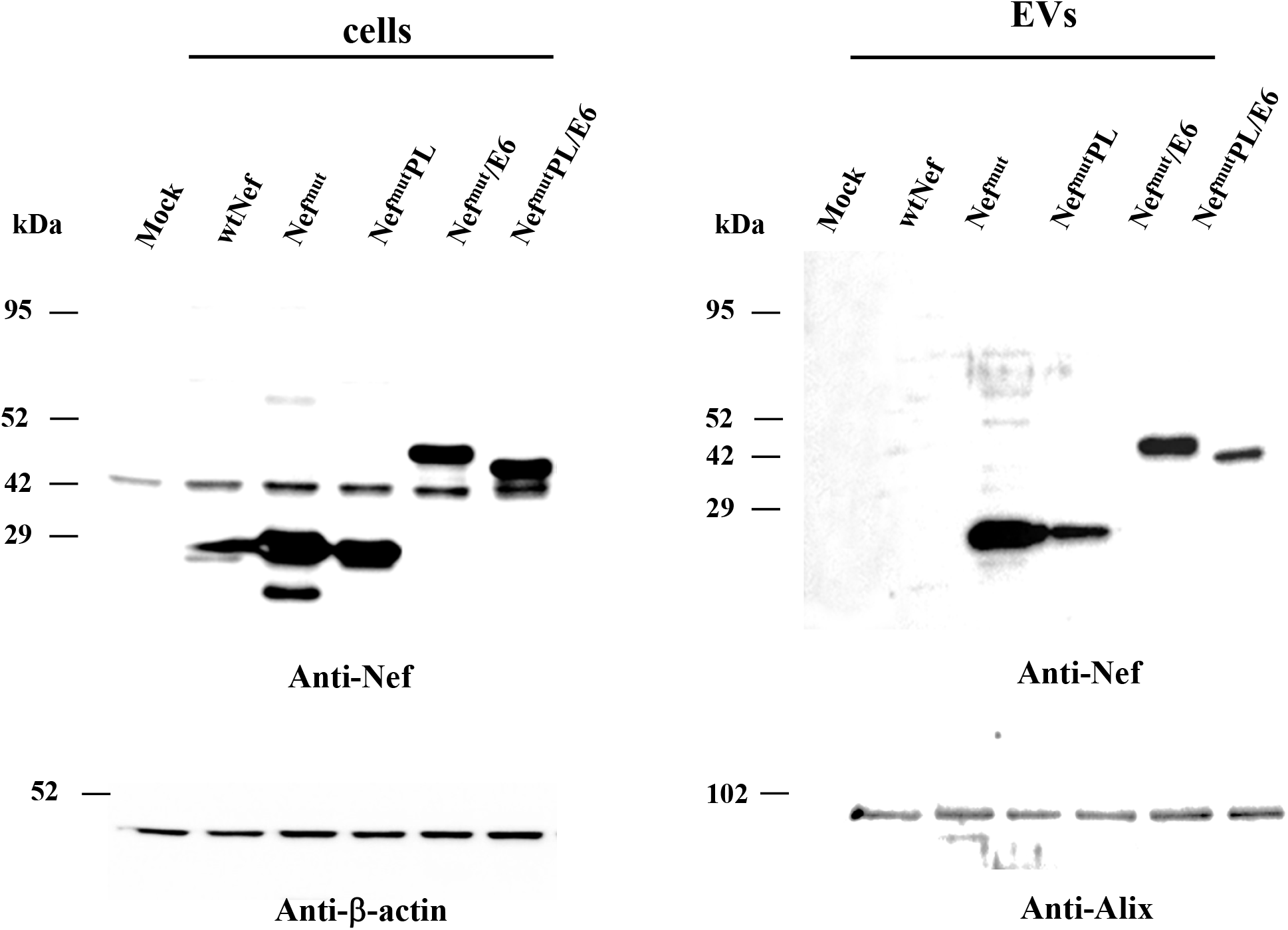
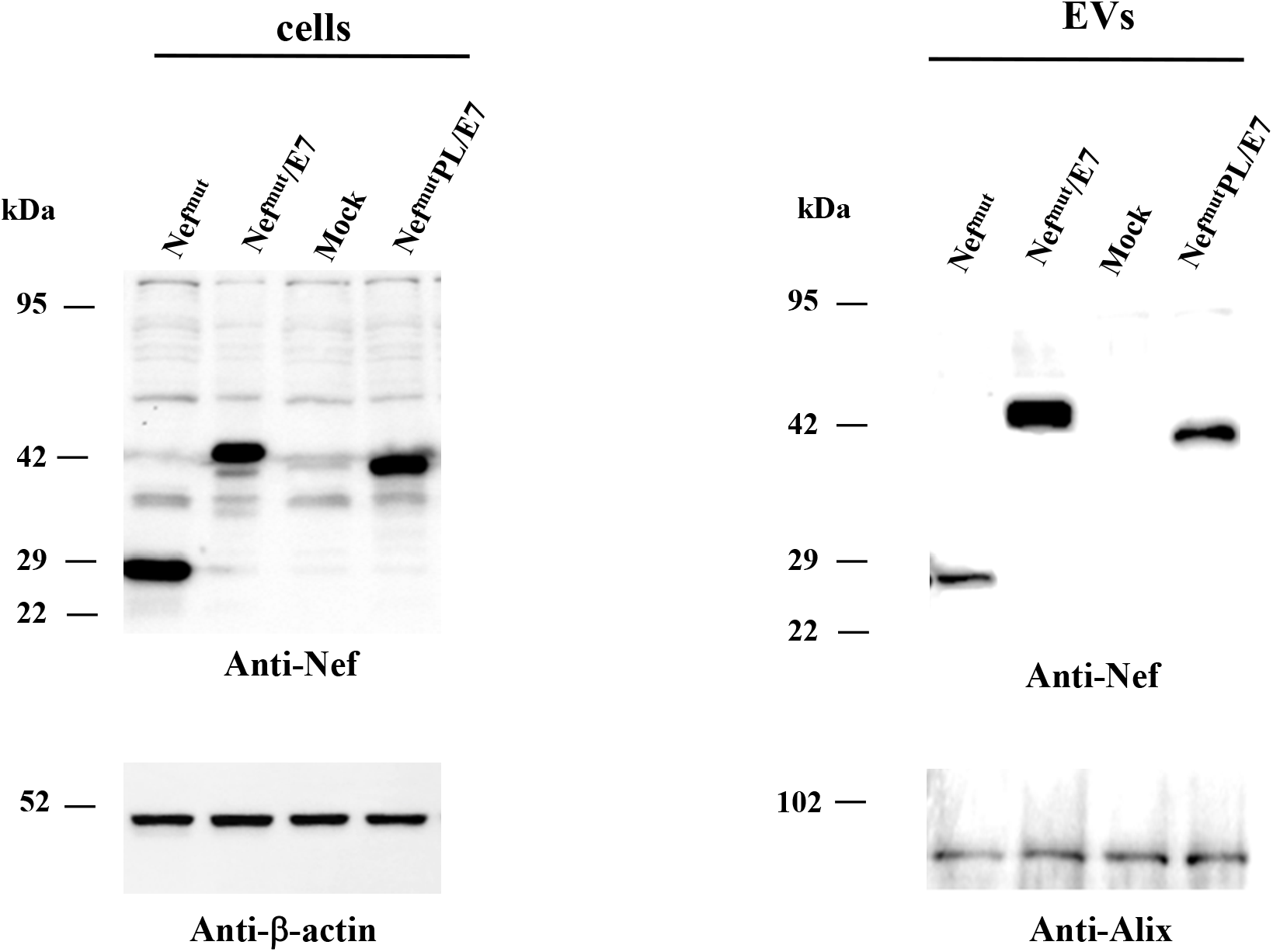
Detection of HPV16-based fusion products in both transfected cells and EVs. A. Western blot analysis on 30 µg of cell lysates from HEK293T cells transfected with DNA vectors expressing the indicated fusion products (left panel), and equal volumes of buffer where purified EVs were resuspended after differential centrifugations of respective supernatants (right panel). As control, conditions from mock-transfected cells as well as cells transfected with either wt Nef, Nef^mut^, or Nef^mut^PL were included. The results are representative of three independent experiments. B. Analysis on 30 µg of cell lysates from HEK293T cells transfected with DNA vectors expressing the indicated fusion products (left panel), and equal volumes of buffer where purified EVs were resuspended after differential centrifugations of respective supernatants (right panel). As control, conditions from mock-transfected cells were included. The results are representative of four independent experiments. For both panels, polyclonal anti-Nef Abs served to detect Nef^mut^-based products, while β-actin and Alix were markers for cell lysates and EVs, respectively. Molecular markers are given in kDa.

We concluded that the presence of the C-terminal 29 amino acids of Nef^mu^ is not essential for the efficient EV-uploading of the HPV16 products fused to it.

### Induction of HPV16-specific polyfunctional CD8^+^ T lymphocytes in mice immunized with DNA vectors expressing Nef^mut^PL/E6 and /E7

The induction of strong CTL immunity through the Nef^mut^-based platform against the heterologous antigen implies the induction of antigen-specific CD8^+^ T lymphocytes co-expressing inflammatory Th-1 cytokines including IFN-γ, IL-2, and TNF-α (i.e., polyfunctional CD8^+^ T lymphocytes). Next, we compared the effectiveness of Nef^mut^ to that of its truncated form in favoring the antigen-specific CD8^+^ T cell immune response against the foreign antigen also in terms of induction of polyfunctional CD8^+^ T lymphocytes.

DNA vectors expressing HPV16-E6 and-E7 fused with either Nef^mut^ or Nef^mut^PL were injected in mice. Fourteen days after the second immunization, IFN-γ EliSpot assays showed that all injected mice responded in terms of antigen-specific CD8^+^ T cell immunity (Fig. 4A). Overall, DNA vectors expressing E7-derivatives induced stronger immune responses compared to E6-derivatives. On the other hand, the levels of CD8^+^ T cell immune response in mice immunized with Nef^mut^PL-based vectors appeared similar to those detected after injection of DNA vectors expressing full-length Nef^mut^ (Fig. 4A). Single-cytokine ICS analysis showed that the percentages of E6/E7-specific CD8^+^ T lymphocytes expressing either IFN-γ, IL-2, or TNF-α did not significantly change in the presence of Nef^mut^ or Nef^mut^PL. As an exception, increased percentages of TNF-α expressing CD8^+^ T cells were observed from mice injected with DNA expressing Nef^mut^PL/E6 compared to those immunized with Nef^mut^ /E6 (Fig. 4B). Most important, the C-terminal truncation of Nef^mut^ did not affect the generation of antigen-specific, polyfunctional CD8^+^ T lymphocytes (fig. 4C).

**Figure 4.**
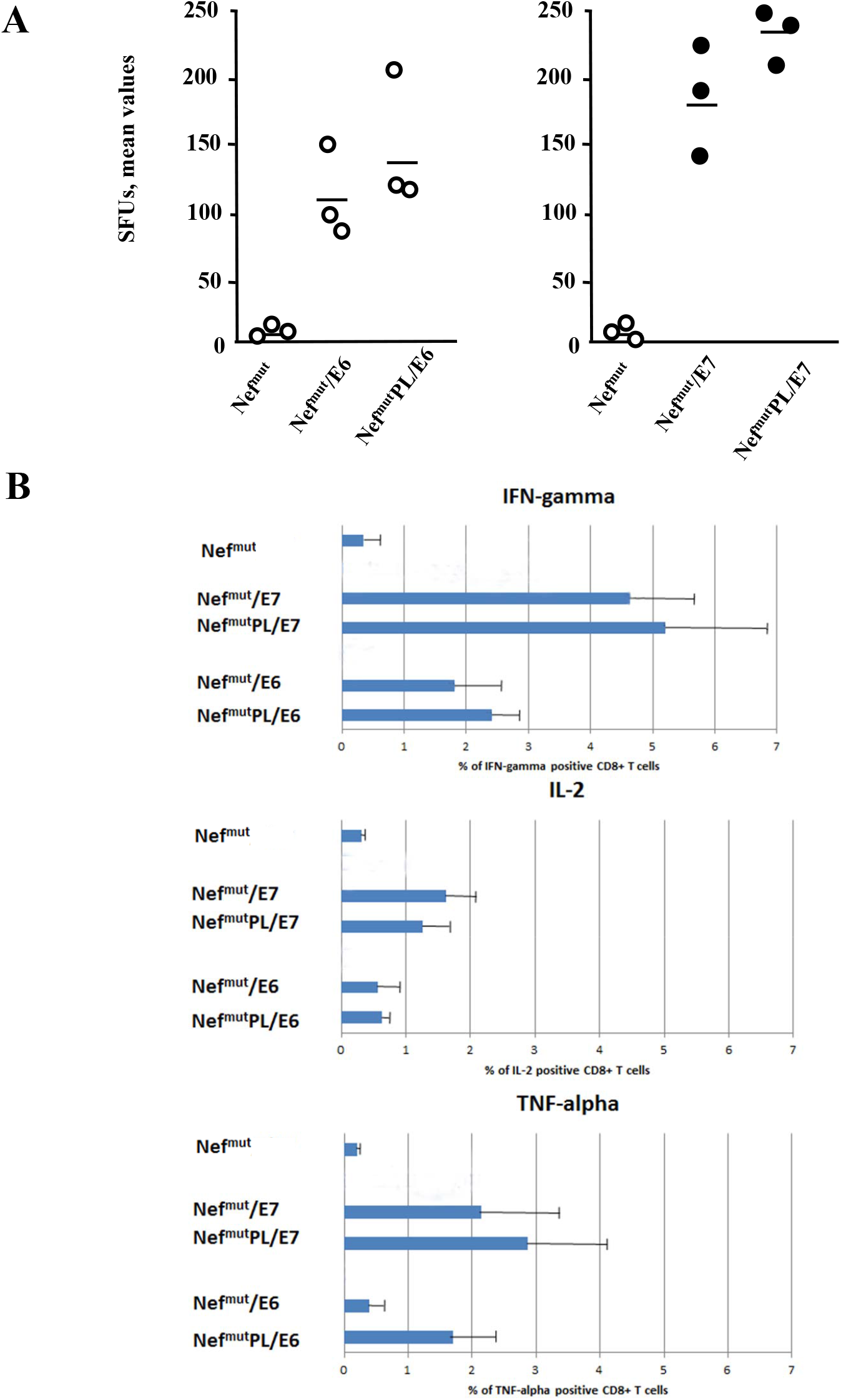
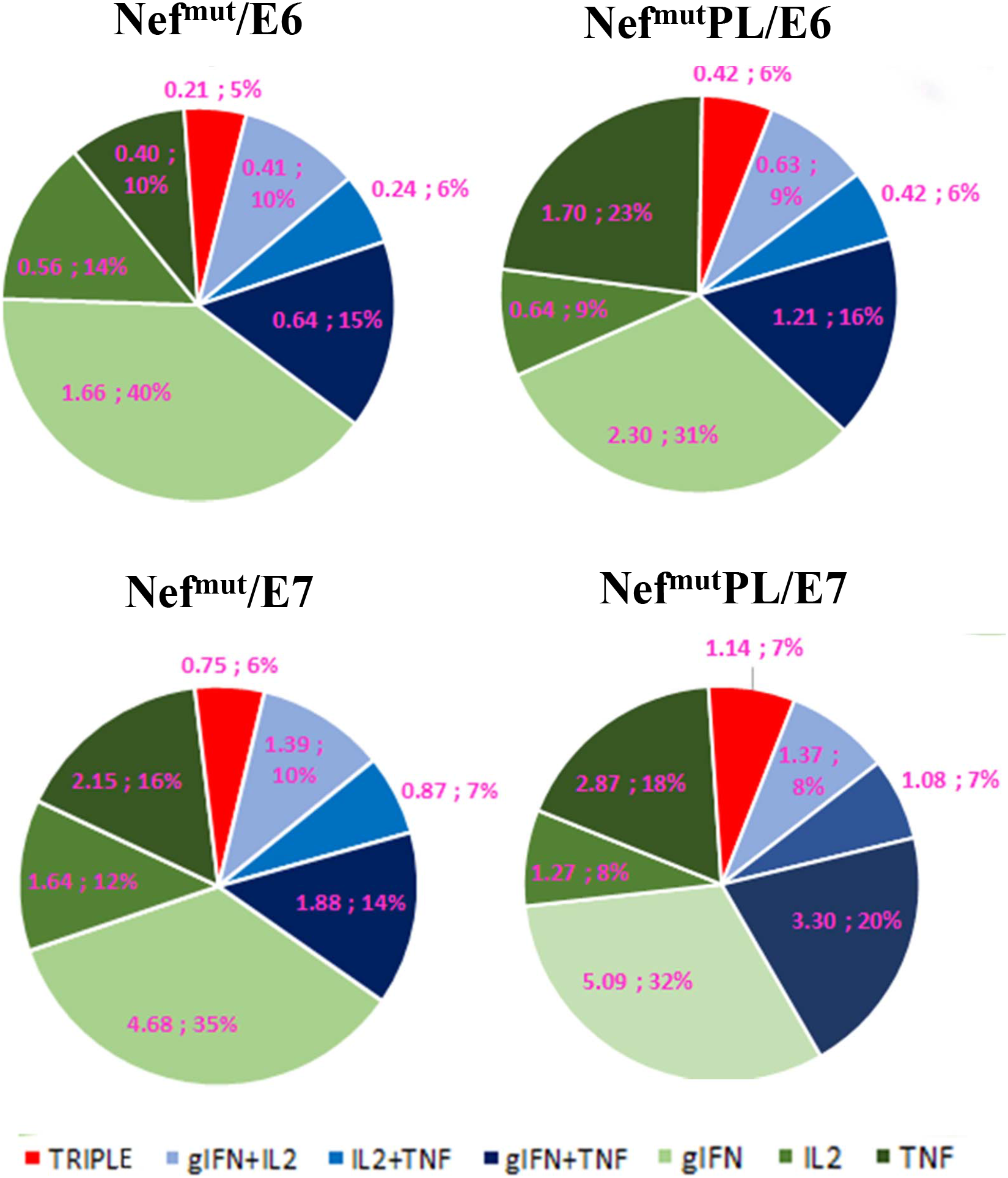
HPV16 E6-and E7-specific CD8^+^ T cell immunity induced in mice after injection of DNA vectors expressing either Nef^mut^ or Nef^mut^PL. A. CD8^+^ T cell immune response in C57 Bl/6 mice inoculated with DNA vectors expressing the indicated fusion products. As control, mice were inoculated with a vector expressing Nef^mut^ alone. At the time of sacrifice, 2.5×10^5^ splenocytes were incubated o.n. with or without 5 µg/ml of either unrelated (not shown), E6, or E7-specific peptides in triplicate IFN-γ EliSpot microwells. Shown are the number of IFN-γ SFUs/well as mean values of triplicates. Intragroup mean values are also reported. B. IFN-γ, IL-2, and TNF-α intracellular accumulation in CD8^+^ T cells from cultures of splenocytes isolated from each mouse injected with the indicated DNA vectors. Splenocytes were incubated o.n. with or without 5 µg/ml of either unrelated, E6 or E7-specific peptides in the presence of brefeldin A, and then analyzed by ICS. Shown are the mean values +SD of positive CD8^+^ T cells within total alive CD8^+^ T cells for each cytokine as calculated after subtraction of values detected in CD8^+^ T cells from cultures treated with the unrelated peptide. C. Pie charts indicating means of both absolute (i.e., over the total of analyzed CD8^+^ T cells) and relative percentages of CD8^+^ T cells expressing cytokine combinations within splenocyte cultures isolated from each mice. Percentages were calculated after subtraction of values detected in homologous cultures treated with unrelated peptides. Values detected with splenocytes from mice injected with control vector were below the sensitivity threshold of the assay. The results are representative of two independent experiments.

We concluded that the effectiveness of the Nef^mut^-based vaccine platform in terms of anti-HPV16-E6 and –E7 immunogenicity remained largely unchanged when the anchoring protein was deleted of its 29 C-terminal amino acids.

### Intracellular expression and EV-uploading of SARS-CoV-2 antigens fused with Nef^mut^PL

Very recently we demonstrated that the Nef^mut^-based vaccine platform could be instrumental to induce strong CD8^+^ T cell immunity against multiple SARS-CoV-2 antigens (22). The immune response was detected in both spleens and lung airways. To establish whether these immunogenic properties were preserved also in the presence of the Nef^mut^ C-terminal truncation, we first evaluated the EV-uploading efficiency of the fusion products. Investigations were carried out with both S1 and S2 SARS-CoV-2 antigens, which were proven to be the most immunogenic antigens in the context of the Nef^mut^ vaccine platform.

Intracellular expression of the fusion products were evaluated by transient transfection in HEK293T cells, and the relative levels of uploading into EVs were scored upon their isolation from the supernatants. Results from western blot assays indicated that S1 and S2 accumulated in cells at comparable extents when fused with the two EV-anchoring proteins (Fig. 5). Most important, no evident differences in the levels of EV-uploading were assessed (Fig. 5).

**Figure 5.**
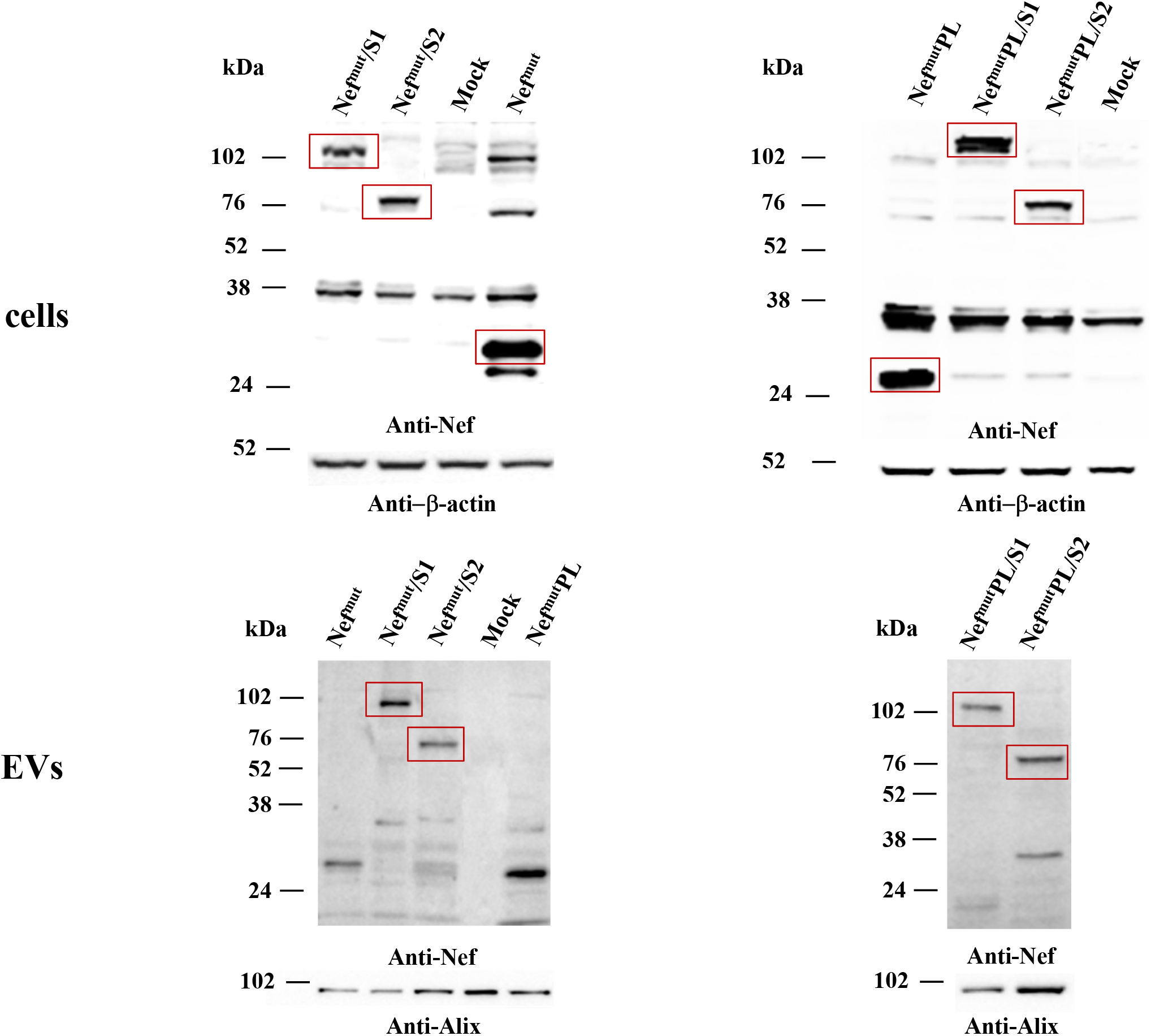
Detection of SARS-CoV-2 S1- and S2-based fusion products in transfected cells and EVs. Western blot analysis on 30 µg of lysates from HEK293T cells transfected with DNA vectors expressing either Nef^mut^ or Nef^mut^PL fused with either SARS-CoV-2 S1 or S2 ORFs (upper panels). Equal volumes of buffer where purified EVs were resuspended after differential centrifugations of the respective supernatants were also analyzed (bottom panels). As control, conditions from mock-transfected cells as well as cells transfected with a vector expressing Nef^mut^ and/or Nef^mut^PL were included. Polyclonal anti-Nef Abs served to detect Nef^mut^-based products, while β-actin and Alix were revealed as markers for cell lysates and EVs, respectively. Relevant signals are highlighted. Molecular markers are given in kDa. The results are representative of two independent experiments.

**Figure 6.**
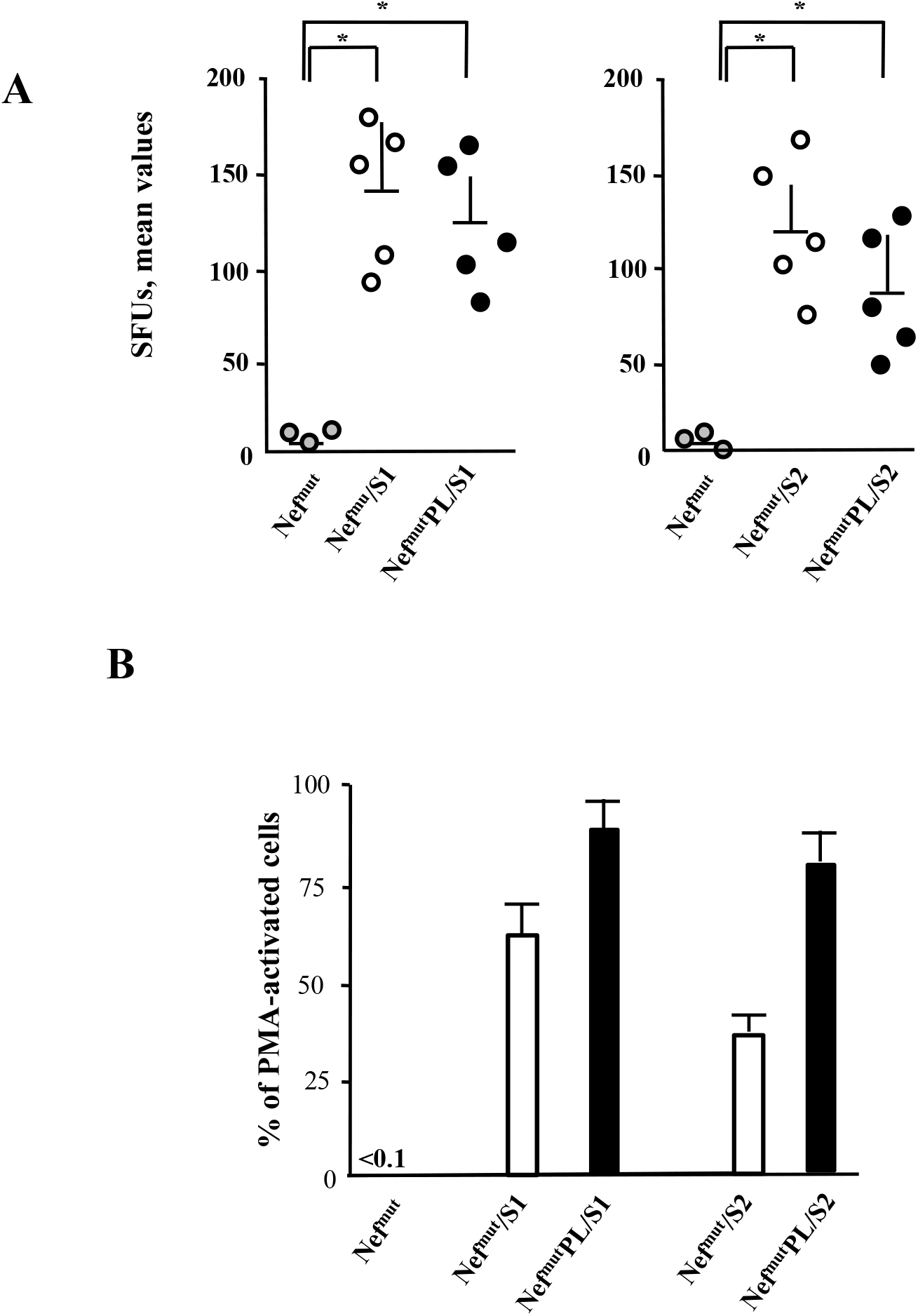
SARS-CoV-2-specific CD8^+^ T cell immunity induced in both spleens and lungs of DNA injected mice. A. CD8^+^ T cell immune response in either C57 Bl/6 (for S1-related DNA) or Balb/c mice (for S2-related DNA) inoculated i.m. with the DNA vectors expressing either Nef^mut^ or Nef^mut^PL fused with the indicated SARS-CoV-2 antigens. As controls, mice were inoculated with a vector expressing Nef^mut^. At the time of sacrifice, 2.5×10^5^ splenocytes were incubated o.n. with or without 5 µg/ml of either unrelated or SARS-CoV-2-specific peptides in triplicate IFN-γ EliSpot microwells. Shown are the numbers of IFN-γ SFUs/well as mean values of triplicates after subtraction of mean values measured in wells of splenocytes treated with the unspecific peptides. Reported are intragroup mean values + standard deviations also. The results are representative of two independent results. Statistical significance compared to values obtained with splenocytes injected with Nef^mut^ expressing vectors was determined by Mann-Whitney U test. ^*^*p*<0.05. B. Percentages of SFUs +SD detected in IFN-γ EliSpot microwells seeded with 10^5^ cells from pooled BALFs in the presence of virus-specific peptides compared to PMA plus ionomycin. Cell samples seeded with unrelated peptides scored at background levels. The results are representative of four independent experiments.

As already observed with HPV16 products, also in the case of SARS-CoV-2 antigens the efficiency of EV uploading was not influenced by the Nef^mut^ C-terminal truncation.

### CD8^+^ T cell immunity induced in both spleen and lungs of mice injected with vectors expressing Nef^mut^PL-S1 and –S2

Both SARS-CoV-2 S1 and S2 were previously shown to induce a strong CD8^+^ T cell immunity in both spleens and lung airways when expressed by a DNA vector as product of fusion with Nef^mut^ (22). Next, we compared the CD8^+^ T cell responses following the injection of DNA vectors expressing the two antigens fused with either Nef^mut^ or Nef^mut^-PL. Fourteen days after the second immunization, splenocytes were isolated from injected mice, and cultured overnight in IFN-γ EliSpot microwells. Comparable antigen-specific CD8^+^ T cell activations were observed in splenocytes from mice inoculated with each vector expressing the diverse fusion products, in the presence of a slight increase with vectors expressing full-length Nef^mut^ (Fig. 7A). On the other hand, the levels of immune responses measured in cells isolated from bronchoalveolar lavage fluids (BALFs) of injected mice appeared similar, in this case with a small increase in cells from mice injected with Nef^mut^PL-based vectors (Fig. 7B).

These data strongly supported the idea that the Nef^mut^ C-terminus is dispensable for the induction of SARS-CoV-2 specific immunity in spleens and, more striking, lung airways, i.e., the peripheral tissue primarily involved in SARS-CoV-2-related pathogenesis. Hence, Nef^mut^PL could be considered as an effective and safe alternative to full-length Nef^mut^ for anti-SARS-CoV-2 CTL vaccines based on the technology of endogenously engineered EVs.

## Discussion

The possibility to translate into the clinic the Nef^mut^-based CTL vaccine platform imposes the need to identify and eliminate unnecessary sequences in both vectors and ORFs coding for the fusion products. Concerning the DNA vector, we proved that Nef^mut^-based fusion antigens maintain unaltered their immunogenic properties when expressed by the pVAX1 vector (22), i.e., a vector where prokaryotic sequences are reduced compared to the more complex pTargeT vector we previously utilized. More advanced, extremely small DNA vectors lacking antibiotic resistance (e.g., minicircle DNA vectors) would represent an interesting alternative for translation into the clinic of the Nef^mut^-based vaccine platform (33). On the other hand, the possibility to reduce the sequences required for the efficient EV-uploading was here explored. Although we largely documented that Nef^mut^ lacks all biologic activities of the wild-type counterpart (3), it should be considered that its defectiveness relies on the co-existence of two amino acid substitutions. Unpredictable, rare, however theoretically possible events of back-mutation might occur, especially when the immunogenic DNA undergoes large-scale production and delivery. In this context, possible anti-cellular effects due to back-mutations are expected to be strongly mitigated by the deletion of a significant part of the protein. In particular, the 29 amino acid C-terminal region of Nef includes domains involved in both signaling and trafficking functions typical of wt Nef (34). In the case of Nef^mut^, we found that the truncation of a C-terminal region of 29 amino acids had no impact on both efficiency of EV uploading and immunogenicity of diverse fusion products. This conclusion can be applied to any foreign antigen, since it was drawn through the analysis of both small- and large-sized proteins from unrelated viruses.

In some instance, our results can be considered unexpected since the Nef C-terminal region comprises a well conserved cholesterol recognizing motif (CRM) already characterized as an important domain for the association with the viral envelope (35). Considering the convergence in the biogenesis of HIV-1 and EVs (36), this domain has the potential to play a role in the interaction of Nef with the nanovesicles released by Nef-expressing cells. The CRM amino acid sequence, i.e., LHPEYYK in SF2-like Nef alleles and LHPEYFK in HXB2-like ones, is well conserved among HIV-1 types B strains, including that from which Nef^mut^ originated, i.e., F12/HIV-1 (37). We hypothesized that the effects of both N-terminal myristoylation and palmitoylation present in Nef^mut^, together with the stretch of basic amino acids located in alpha helix 1 are prevalent over those induced by CRM in terms of stable interaction with lipid rafts of both plasma and endosomal membranes.

Overall, the possibility to include Nef^mut^PL sequences in DNA vectors candidates for new vaccine strategies against infectious and tumor diseases significantly strengthens the safety profile of the EV-based CTL vaccine platform.

## Ackowledgements

We thank Antonio Di Virgilio and Pietro Arciero, Istituto Superiore di Sanità, for technical support.

